## Supplementary Text for "Increased antibiotic susceptibility in *Neisseria gonorrhoeae* through adaptation to the cervical environment"

Count tables and statistical analysis for exploratory dataset (global meta-analysis):

```
"Overexpression background and mtrC LOF"
  upregulation mtrC_prematureStop     n
  <chr>        <chr>              <int>
1 no           Intact               617
2 no           LOF                    6
3 yes          Intact              4028
4 yes          LOF                  174
5 NA           Intact                52
6 NA           LOF                    5


"Site of infection and mtrC LOF"
       Cervix Pharynx Rectum Urethra
Intact    113     103    242    2167
LOF        16       3      4      82

Fisher's Exact Test for Count Data

data:  site.chisq
p-value = 6.487e-05
alternative hypothesis: two.sided


"Sexor and mtrC LOF"
       WSM MSW MSMW  MSM
Intact  30 598  123 1158
LOF      3  28    4   31

Fisher's Exact Test for Count Data

data:  sexor.chisq
p-value = 0.04021
alternative hypothesis: two.sided
```

```
"Overexpression background and mtrA LOF"
  upregulation mtrA            n
  <chr>        <chr>       <int>
1 no           full_length   355
2 no           LOF           268
3 yes          full_length  4113
4 yes          LOF            89
5 NA           full_length    52
6 NA           LOF             5


"Site of infection and mtrA LOF with no mtr upregulation"
          Cervix Pharynx Rectum Urethra
Inducible    104     102    242    2187
LOF           25       4      4      61

Fisher's Exact Test for Count Data

data:  site.chisq
p-value = 1.636e-12
alternative hypothesis: two.sided


"Sexor and mtrA with no mtr upregulation"
          WSM MSW MSMW  MSM
Inducible  27 598  127 1181
LOF         6  28    0    7

Fisher's Exact Test for Count Data

data:  network.chisq
p-value = 1.805e-11
alternative hypothesis: two.sided
```

```
"Site of infection and farA LOF"
       Cervix Pharynx Rectum Urethra
Intact     96     103    234    2129
LOF        33       3     12     117

	Fisher's Exact Test for Count Data

data:  site.chisq
p-value = 1.775e-12
alternative hypothesis: two.sided


"Sexor and farA LOF"
       WSM MSW MSMW  MSM
Intact  25 585  125 1166
LOF      8  40    2   22

	Fisher's Exact Test for Count Data

data:  sexor.chisq
p-value = 5.055e-10
alternative hypothesis: two.sided
```

Count tables and statistical analysis for validation dataset:

```
"Overexpression background and mtrC LOF"
  upregulation mtrC_prematureStop     n
  <chr>        <chr>              <int>
1 no           Intact                89
2 no           LOF                    2
3 yes          Intact              2060
4 yes          LOF                   33
5 NA           Intact                 2


"Overexpression background and mtrA LOF"
  upregulation mtrA          n
  <chr>        <chr>     <int>
1 no           11-bp Del     4
2 no           Intact       87
3 yes          11-bp Del    81
4 yes          Intact     2012
5 NA           Intact        2


"Site of infection and mtrC LOF"
       Cervix Pharynx Rectum Urethra
Intact    218     383    625     867
LOF         9       3      7      15

Fisher's Exact Test for Count Data

data:  site.chisq
p-value = 0.02561
alternative hypothesis: two.sided


"Sexor and mtrC LOF"
       WSM MSW  MSM
Intact 268 243 1424
LOF     10   4   17

Fisher's Exact Test for Count Data

data:  sexor.chisq
p-value = 0.01803
alternative hypothesis: two.sided
```

```
"Site of infection and farA LOF"
       Cervix Pharynx Rectum Urethra
Intact    160     354    609     782
LOF        67      31     21      98

	Fisher's Exact Test for Count Data

data:  site.chisq
p-value < 2.2e-16
alternative hypothesis: two.sided


"Sexor and farA LOF"
       WSM MSW  MSM
Intact 197 186 1386
LOF     81  60   51

	Fisher's Exact Test for Count Data

data:  sexor.chisq
p-value < 2.2e-16
alternative hypothesis: two.sided
```
